# Stimulus Onset Hub: An open-source, low latency, and opto-isolated trigger box for neuroscientific research replicability and beyond

**DOI:** 10.1101/721803

**Authors:** Charles E. Davis, Jacob G. Martin, Simon J. Thorpe

## Abstract

There is currently a replication crisis in many fields of neuroscience and psychology, with some estimates claiming up to 64% of research in psychological science is not reproducible. Three common culprits which have been suspected to cause the failure to replicate such studies are small sample sizes, “hypothesizing after the results are known,” and “p-hacking.” Here, we introduce accurate stimulus timing as an additional possibility. Accurate stimulus onset timing is critical to almost all psychophysical research. Auditory, visual, or manual response time stimulus onsets are typically sent through wires to various machines that record data such as: eye gaze positions, electroencephalography, stereo electroencephalography, and electrocorticography. These stimulus onsets are collated and analyzed according to experimental condition. If there is variability in the temporal accuracy of the delivery of these onsets to external systems, the quality of the resulting data and scientific analyses will degrade. Here, we describe an approximately $200 Arduino based system and associated open-source codebase which achieved a 5.34 microsecond delay from the inputs to the outputs while electrically opto-isolating the connected external systems. Using an oscilloscope, the device is configurable for different environmental conditions particular to each laboratory (e.g. light sensor type, screen type, speaker type, stimulus type, temperature, etc). This low-cost open-source project delivered electrically isolated stimulus onset Transistor-Transistor Logic triggers with a median precision of 5.34 microseconds and was successfully tested with 7 different external systems that record eye and neurological data.

## Introduction

The replication crisis in cognitive neuroscience studies poses a substantial challenge for scientists (1–18). Current trends to address the replication crisis suggest several courses of action. First, pre-registration has been suggested in order to eliminate p-hacking and hypothesizing after the results are known. Second, larger sample sizes and better statistics will provide better confidence in results. In addition, opening up access to data and analysis code will provide ways to find mistakes in analyses that may have been the source of the replication crisis.

Here, we explore accurate stimulus timing as an additional component of the replication crisis. As a potential solution to the complex problem of detecting analog and digital events and delivering them to different systems without significant delay, we describe an inexpensive and ultra-low latency device that we designed and built to detect visual and auditory stimulus onsets and deliver them to multiple external data acquisition systems in an opto-isolated manner. The code and design of the system, with all its parts, along with suggested computer hardware for psychophysical experiments are further described at the project’s website (https://stimulusonsethub.github.io/StimulusOnsetHub/).

Multimodal experimentation in neuroscience is becoming common practice. Not only can inputs be multimodal (e.g. auditory stimuli, visual stimuli, somatosensory stimuli, *etc.*), but the output devices to which these input triggers are sent are also potentially multimodal (e.g. EEG+eyetracking, EEG+fMRI, EcoG+SEEG+eyetracking, *etc.*). Multimodal recordings from multimodal inputs pose technical challenges that relate to electrical interference, crosstalk, and temporal precision of stimulus onsets. The experiments in our neuroscience lab generally explored humans’ capability to do ultra-fast continuous face and sound detection (19–21). The experiments operated at high speeds, and we needed to align the visual and auditory stimulus onsets with various combinations of recording equipment such as eye tracking, electroencephalography (EEG), stereo electroencephalography (SEEG), and electrocorticography (ECoG) (22). The delay for detecting and delegating the recorded triggers to external equipment was an important factor (23). It is tolerable to have such delays as long as they can be accounted for accurately and predictably. Ideally, however, such delays should be kept to a minimum. After experimentation and optimization, the input/output lag of the device had a median precision of 5.34 microseconds (see Figure 5).

In addition, as these recording systems are sometimes sensitive to electrical crosstalk, we designed the device to opto-isolate the different recording systems from each other. When multiple recording systems are connected at the source of the stimulus detector, ground loops between the systems can create unwanted noise. As a solution, opto-isolation of these different recording systems from each other can reduce noise and thereby improve data quality.

We requested quotes from 3 different companies and received prices ranging from 1,645 to 4,488 US dollars. Each company offered add-ons to interface with more than one device, but these were also rather expensive. One company offered an adapter for our BioSemi Active Two (BioSemi, Amsterdam, Netherlands) data acquisition system for 125 US dollars and one for our Eyelink 1000+ (SR Research, Ottawa, ON, Canada) eye tracker for 55 US dollars. Another company charged 395 US dollars for all 3 connections that we needed. No company seemed to offer opto-isolation, but we couldn’t be sure without opening each device. Instead, we spent about 200 US dollars and made the opto-isolated device ourselves, along with an open-source Arduino codebase.

Our task was to design and build a robust device that would perform its duties with minimal delay and also minimize electrical crosstalk between the different connected system. We were concerned about potential data quality loss associated with noise coming into our EEG, SEEG, ECoG, and eye tracking systems due to potential ground loops and crosstalk created at the different connections to the trigger hub. In order to alleviate these concerns we used opto-isolators to create electrically decoupled barriers between the inputs and the outputs of our device. Next, the device interfaced with the photodiode and microphone recording devices and the experimental display screen in order to accurately time stimulus onsets within the experiments (see Figure 1). The light and sound detectors bypassed any unknown additional latencies introduced by the stimulus computer, operating system, experimental application, graphics card, sound card, speakers, etc (see Figure 2) (25). Finally, we wanted to be able to control multiple trigger propagation, trigger thresholds, and input/output times in order to prevent unaccounted for triggers and inconsistent usage.

**Fig. 1.**
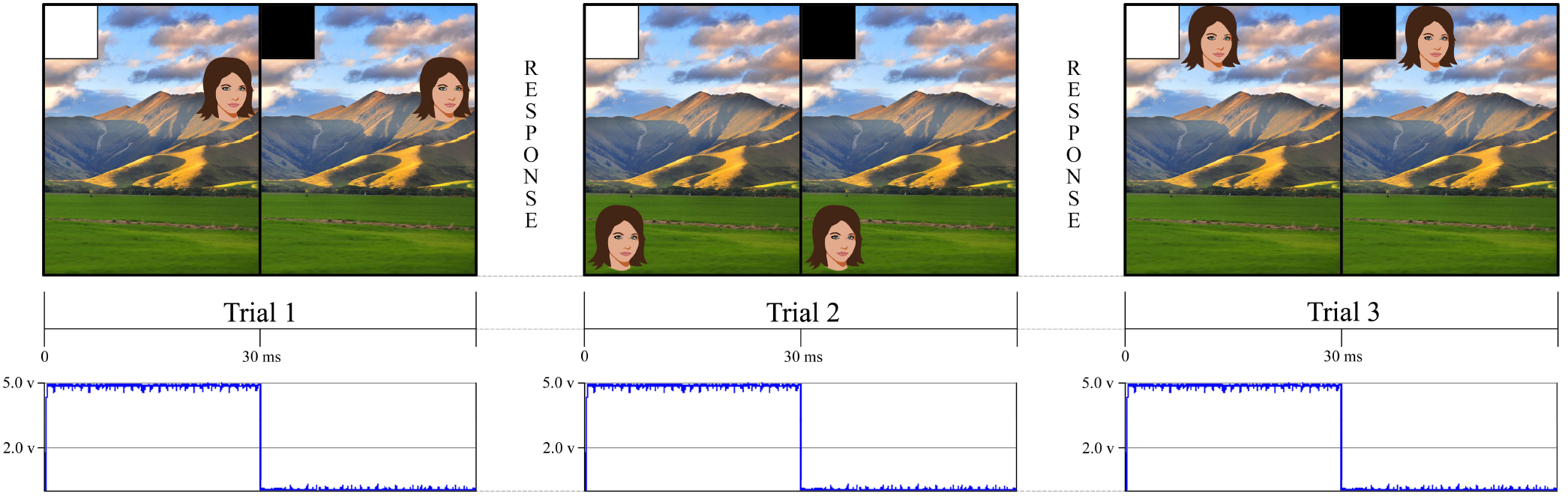
This figure is an example of a simple experiment using an eye tracker and a stimulus computer. The eye tracker follows the eyes as the they saccade and affix on face targets. The stimulus screen presents a white square in the upper left hand corner for 30 milliseconds to indicate when the trial has begun and then a black square, in the same location, until the stimulus system is ready to present another trial. Our stimulus tracking device used a photodiode affixed to the screen to detect when the white square appeared. This allowed us to independently verify when an event happened rather than relying on the stimulus presentation system to tell us when the event happened. For landscape photo attribution see (24).

**Fig. 2.**
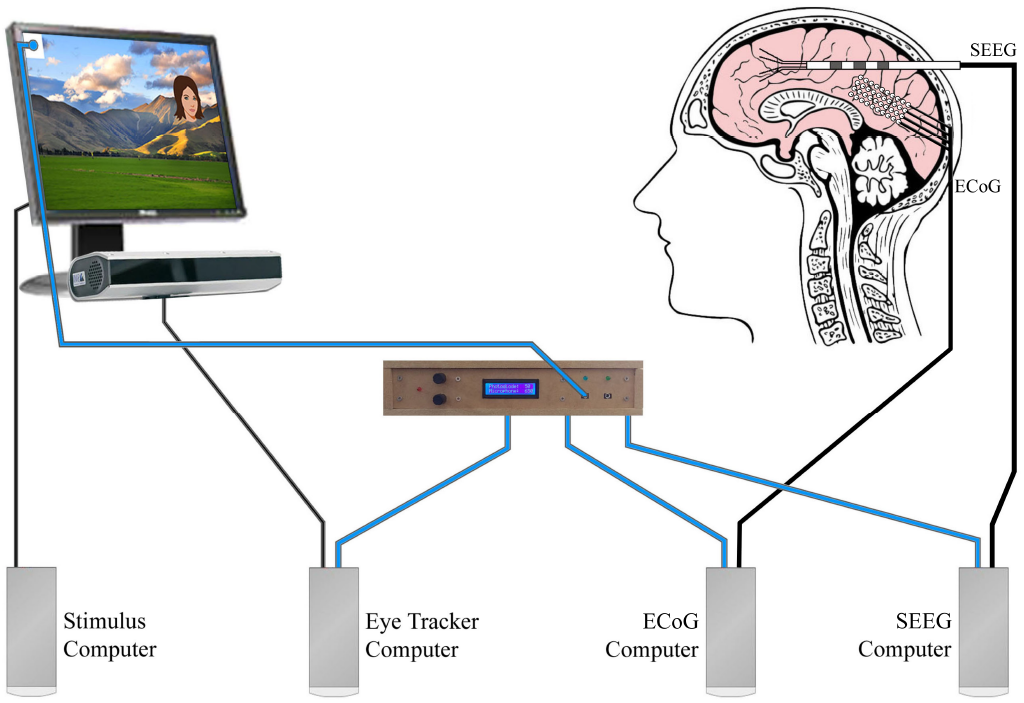
This figure uses the same experiment from Figure 1. This is a simple multimodal experiment where we demonstrate how our device bypasses all systems to provide a low latency onset stimulus to any number of external recording devices. Blue wires carried the photodiode signal to the external recording systems. This multidevice experiment used eye tracking, ECoG, and SEEG. Decoupling the systems with opto-isolators prevented potential interference that could arise due to differential grounding and electrode proximity. For landscape photo attribution see (24).

We used the Arduino platform, which has previously been used in a variety of laboratory based experiments for behavioral research (26) and neural research (27). We tested and optimized the input and output lag according to our particular experimental conditions and hardware. The project ultimately saved us thousands of dollars over other commercial systems that were not opto-isolated, while also improving performance. The device, in conjunction with an oscilloscope, allowed us to find additional latencies introduced by other experimental devices and improved the quality of our data by eliminating electrical crosstalk and ground loops between external connected devices.

## Materials

Here we describe our design and why we made different design decisions. We describe in detail how we realised and built this device with circuit diagrams, Arduino pin maps, benchmarking code, and Arduino C code.

### Device Design

The Arduino Mega 2560 Rev3 is an open-source microcontroller board based on the ATmega2560 microcontroller. We chose the Arduino Mega because it offered the most ports in the Arduino family. It has 54 digital input/output pins with 14 of these pins available for use as pulse width modulation (PWM) outputs, 16 analog inputs, a 16 MHz crystal oscillator, a USB connection, and a power jack. The Arduino can be powered by a computer or a USB cable, but, according to the documentation, if supplied with less than 7 volts, the 5 volt pin may supply less than 5 volts, and the board may become unstable. We found this to be true when we tested our device. To overcome this problem, we chose a 12 volt, 1 amp AC-to-DC adapter (wall-wart) with a 2.1 mm center-positive barrel connector.

Breakout boards with screw terminal blocks were chosen for wired connections in order to provide a more robust development platform for easy prototyping and hassle free testing with an oscilloscope. We wanted to avoid soldering, as well as desoldering, as much as possible. We also wanted to avoid having to buy expensive crimping tools to make connections such as Dupont and Japan Solderless Terminal (JST) wire connections. Additionally, WAGO brand splicers were implemented for further ease of use. We used multicolored, 22 American Wire Gauge (AWG) wire from Adafruit Industries that was the exact gauge of the Arduino ports. We found that the wires with Dupont connectors that came with most Arduino starter kits did not feel secure in the Arduino Mega ports, but the Adafruit wires definitely seated better.

In order to confront the problems of noise due to electrical crosstalk and ground loops, a dual channel, opto-isolator break out board made by SparkFun Electronics was chosen in order to isolate the input side of our stimulus tracker from the output side. It is important to note that, since the inputs were isolated from the outputs, the output side would need a power supply. Fortunately, all but one of the recording devices we tested for compatibility provided several power and ground pins. This recording device did not provide a power source because it had only BNC input connections. To work around this problem, we used a battery pack with an on/off switch which could produce approximately 4.5 volts for running the output side of the opto-isolator for the BNCs. It was necessary to solder screw terminal blocks for these break out boards.

A 1602 liquid-crystal display (LCD) screen with an I2C bus from SunFounder was incorporated into the user interface. In order to display the input device sensitivity levels that were used to control trigger sensitivity, we picked a LCD with a pre-attached I2C bus which helped reduce wired connections to the Arduino. Two rotary encoder break out boards with plastic knobs were used to control input device threshold levels. For indicator lights, we used red, blue, and green LEDs. Soldering was required to attach the LEDs to their respective 100 ohm resistors and wires. We used audio jack break out boards with pre-attached screw terminal blocks from Gravitech Electronic Experimental Solutions and screw terminal block to male 3.5 mm audio connectors for wiring the photodiode and connecting to the audio jacks. The photodiode was made from version two of an analog ambient light sensor from DFRobot due to its published light ramp up time of 15 microseconds (28).

On the output side of the device, D-subminiature (DB) 25 and 37 breakout boards were purchased from CZH-LABS with pre-attached screw terminal blocks. We used electromagnetically shielded DB25 and DB37 cables with adapters to connect the outputs to the different recording systems. We chose Bayonet Neill–Concelman (BNC) connector breakout boards, also from Gravitech Electronic Experimental Solutions, with pre-attached screw terminal blocks.

### Code Design

We programmed the hardware in the Arduino C Language and made it freely available on the Internet and named it StimulusOnsetHub (29). On the GitHUB page (https://stimulusonsethub.github.io/StimulusOnsetHub/), there is also a hardware list and more detailed information for how to build the device along with suggested tools, computer hardware, and other components for building highly accurate psychophysical experiments. In the code, the "setup()" function is called once when the Arduino powers up, and sets up all pin modes and interrupts for the different connections. The "loop()" function is called repeatedly while the box is running and handles:

- Reading the photodiode and microphone input levels,
- Sending output signals if the input levels cross the user-specified threshold level for each input, and
- Optionally updating the LCD if an interrupt from one of the rotary encoders was triggered.

The performance of the “loop()” function largely determined the temporal delay between the inputs and outputs of the box, aside from any external delays caused by other factors. Interrupts were used for the rotary encoders to avoid the performance reduction that would occur if the rotary knobs’ positions were read each time during the loop. Aside from this early optimization, we tried to avoid other premature optimizations: “...the root of all evil (or at least most of it) in programming (30).” That is, at the end of the software and hardware design, we built compiler switches in the code to test the effects of different combinations of optimization choices with the oscilloscope. We explored several different on/off binary compiler options, such as:

- LIGHTSENABLED: Optionally turn off the LCD and LED lights of the box.
- DIGITALWRITEFAST: Optionally use the digital-WriteFast library (31).
- DIGITALMICROPHONE Optionally use a digital microphone.
- ADDA: Optionally support the analog daughterboard.
- FASTADC: Optionally set different pre-scale methods on potentially connected analog to digital converters.
- DEBUG: Optionally enable debug statements to be sent to the console.

We encountered a limitation with the Arduino in that we could only use one of the analog inputs at a time because of pre-existing time constraints on switching between the reading of different analog channels. The Arduino could only check the analog input rail one analog input at a time. For example, if a photodiode was hooked up to analog input A0 and an analog microphone was hooked up to analog input A1, the Arduino could not read the signal from A0 and the signal from A1 at the same time. Instead, there was a hardware determined delay between the reading of two different analog channels. Thus, the code includes a second option (ADDA) to allow the use of a separate analog to digital daughter board. This setup allowed us to control the threshold of an analog microphone, and was faster than reading from A0 and then switching to A1, but still included a hit in performance as the code in "loop()" had to do more work. Ultimately, we decided to use a digital microphone with an integrated potentiometer that was turned to the most sensitive setting which still reliably detected sound. All wiring diagrams shown in this paper correspond to using an analog photodiode and a digital microphone (i.e. Figures 3 and 4).

**Fig. 3.**
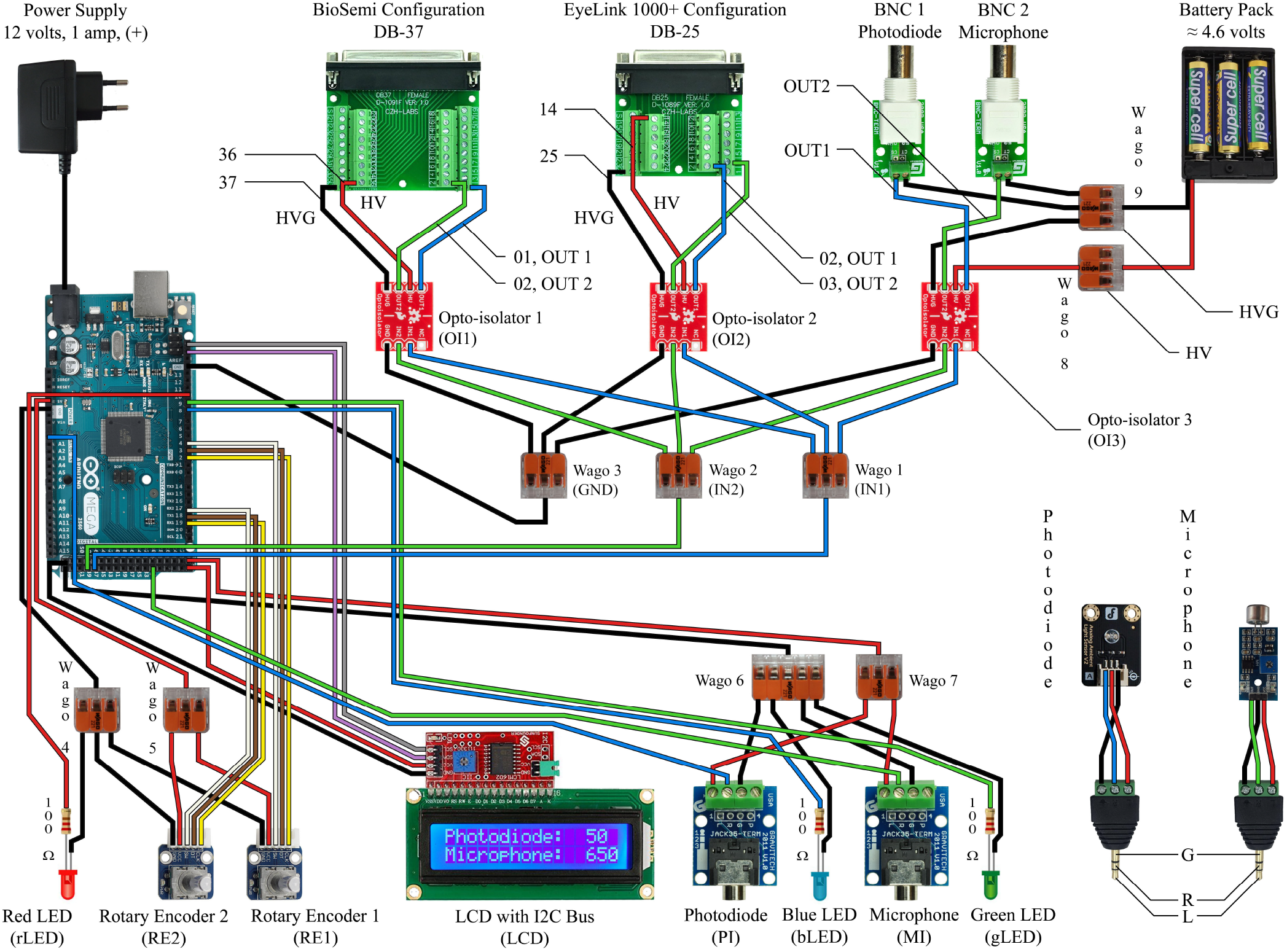
This circuit diagram describes the device we made to work with our EEG system and eye tracker, and it is to be read in conjunction with the Arduino Mega 2560 Rev3 Pin Map (reference Figure 4). The red and black wires correspond to power and ground. The white, brown, and yellow wires are only for the rotary encoders. The gray and violet wires correspond to the SCL and SDA connections for the LCD. The blue wires indicate the path of the photodiode input signal to the output signal as well as the blue indicator light. The green wires indicate the path of the microphone input signal to the output signal as well as the green indicator light.

**Fig. 4.**
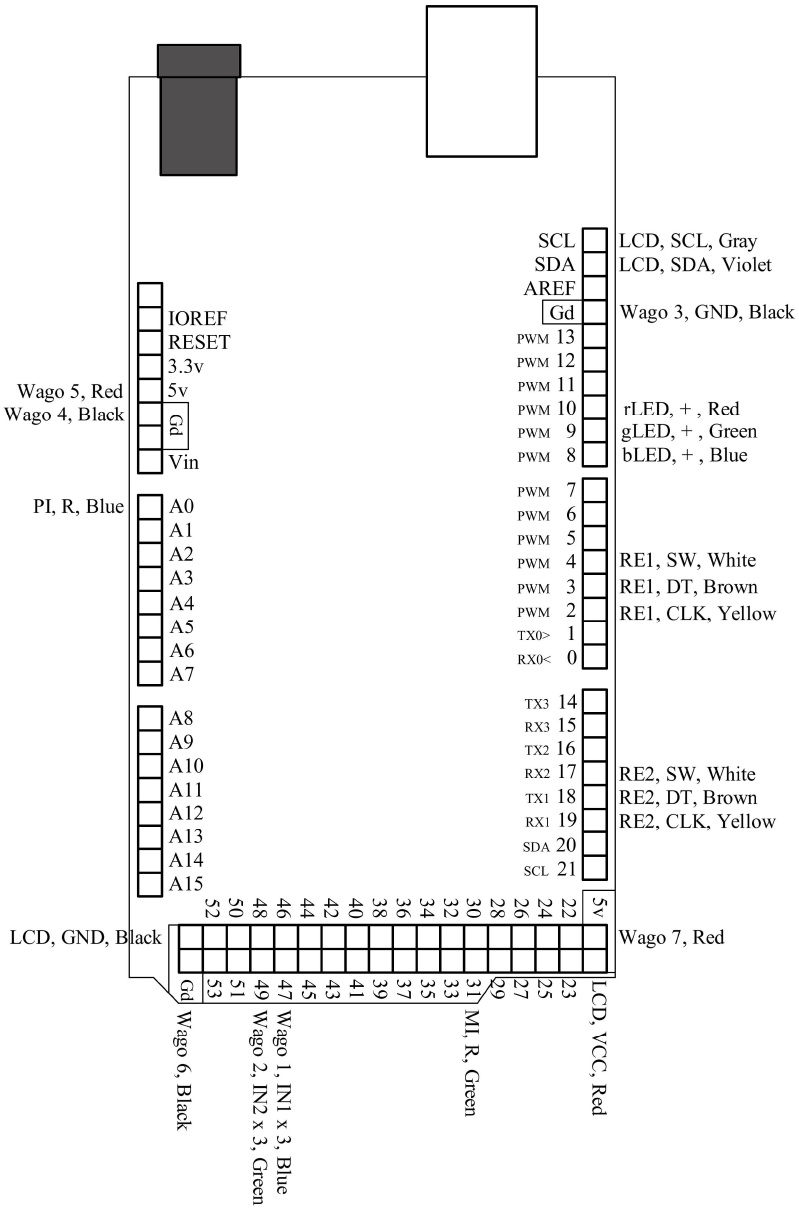
This is a pin map for the Arduino Mega 2560 Rev3. All used ports are labeled and correspond to their respective connections in Figure 3.

## Results

We wrote a benchmarking tool in Matlab using Psychtoolbox (32). The Psychtoolbox program alternately painted 100ms black and white squares on an ASUS PG278Q computer screen to rapidly trigger the photodiode. This LCD computer screen has been shown to give accurate results (3). After launching the benchmarking tool, it cycled through 252 on/off blips. We then used the oscilloscope to measure the latency between the rising edges of the input signals and the rising edges of the output signals.

We used the front 3.5 mm audio port to track the photodiode input signal and the DB25 connector on the back of the device to track the output signal. We placed a short piece of wire in the screw terminal block port for the photodiode input signal and another short piece of wire into the ground connection on the Arduino. Next, we inserted short pieces of wire into the screw terminal block ports of the DB25 for the output signal and the ground. On the oscilloscope, 1 hook-tip probe was attached onto the channel 1 connection and another onto the channel 2 connection. In order to set up this experiment, we first placed the hook-tip probe for channel 1 on the testing point on the photodiode screw terminal block and the alligator clip ground to the ground connection on the Arduino. To acquire the output signal, we placed the hook-tip probe for channel 2 on the signal testing point on the DB25 screw terminal block and the alligator clip ground to the ground connection on the DB25 screw terminal block. It was necessary to hook up the Eyelink eye tracking system to the DB25 connection in order to provide power for the output side of the opto-isolator.

All timing delays through the box were computed using the Tektronix TBS1064 digital oscilloscope. Before each experiment, we allowed the oscilloscope to warm up for 20 minutes as suggested by the user manual. We also made sure to perform low-frequency compensation on all probes prior to testing. Additionally, we used the Do Self Calibration utility, which was provided in the oscilloscope Utility menu, to account for any temperature variation that may have occurred in our testing environment since the last calibration. On the oscilloscope, we set the voltage scale for the photodiode input to 1 volt and the time scale to 10 microseconds. In order to compare the two signals, the vertical position of channel 2 (the TTL output after the opto-isolator) was moved on top of channel 1 (the analog photodiode input), and the oscilloscope was set to averaging mode. Channel 2 was used as the triggering line, the trigger level was put at 3 volts, and the oscilloscope was set to detect the rising edge of the signal.

As verified with the oscilloscope, the latency through the box had a median input/output delay of 5.34 microseconds. The photodiode we used had a reported delay of 15 microseconds. Thus, the total latency to detect and transmit an opto-isolated signal to multiple devices was around 20 microseconds. We thought about trying for even faster delays. However, the Arduino Mega runs at 16 MHz, which corresponds to a clock cycle of 62.5 nanoseconds. Each line of code is broken into several assembly instructions that each take 2-3 cycles or more. Nevertheless, the accuracy we achieved was more than adequate for our purposes because sampling rates for these neural and eye tracking systems are on the order of 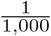 to 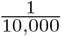 seconds, which is respectively 187.3 to 18.73 times slower than a 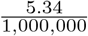 second delay. So, even if we had achieved a better precision, it wouldn’t have helped with the neural and eye tracking technologies that were currently available.

We started our compatibility tests with the Eyelink eye tracker system and then with the BioSemi EEG recording system, but our device will work with any device that can receive a TTL input. Additionally, we had access to other devices to test for compatibility, which included the Red and IViewX line of eye trackers from SMI (SensoMotoric Instruments GmbH, Teltow, Germany), and the Blackrock (Blackrock Microsystems, Inc., Salt Lake City, UT, USA), Neuralynx (Neuralynx, Inc., Boseman, MT, USA), and Micromed (Micromed SAS, Lyon, France) lines of brain recording devices. One drawback of the do it your self approach was that a given design was not completely plug and play across the 7 different recording systems because each had idiosyncratic connection types. Thus, depending on the desired device with which to interface, sometimes it was necessary to spend time going through user manuals, pin out schematics, and internet forums to find out how to make each individual device work with this design. Other times, a voltmeter was needed to map out every pin connection on a device. However, these are unavoidable realities of interfacing signals with different types of modern hardware.

The design should be used in conjunction with an oscilloscope during calibration because experimental conditions such as the particular sensors used and even the environmental temperature can affect sensor readings and sensitivities (33). The more precisely the threshold levels were set for the inputs to the Arduino device, the lower the latency between input and output signals through the device. This occurred because the photodiode input was an analog signal. Therefore, one functional improvement would be to provide an internal oscilloscope or an automatic input/output delay calculation function to tune the device according to the input level thresholds and input/output delay compensations. One could splice the signal output from the photodiode input two ways from within the box. One direction would go to the Arduino as normal, and the other would go to an additional BNC breakout board on the front of the device for hooking up one channel of an oscilloscope. A poor splice can affect both the voltage and current levels. Thus, the best photodiode threshold level might change when splicing the photodiode towards both the oscilloscope and the Arduino. To solve this problem, one could just leave the oscilloscope connected for the experiments to be sure of the timing delay and the best settings for the input thresholds. Therefore, one could essentially have an online oscilloscope during each experiment to store verification data for the trigger timing for each trial. The length of the wire for the photodiode also can contribute a very small amount of delay between photodiode onset and photodiode analog delivery time. So, perhaps the conductivity and temporal delay from the photodiode to the box should also be accounted for with an oscilloscope.

## Discussion

The replication crisis in cognitive neuroscience inspired us to investigate all sources of delay and noise in our experiments so that our efforts might be better replicated (1–18). If 64% of all psychological science is not reproducible, what percent of this 64% is due to errant stimulus timing or unshielded systems? To ensure that our results were not affected by these potential problems, we built a device for detecting and delivering visual and audio events to multiple systems.

Combinations of different types of stimuli (auditory, visual, manual reactions) and neural recording devices complicated the coordination of sending stimulus onsets to different systems both electrically and temporally. When multiple systems are connected at the source of the stimulus processor, electrical interference can occur between systems, leading to undesirable crosstalk which can degrade the quality of the data. Commercial products exist to handle this problem, but we were unable to find any that electrically isolate the connected systems or that perform with the ultra-fast processing that we measured in the device that we built. In addition, with the use of opto-isolators, the quality of the data was improved. The DIY aspect of this project gave us more fine tuned control. We also verified first hand how our devices performed while saving thousands of dollars which was instead put toward other aspects of the overall research project. Although this device was designed with a neuroscientific environment in mind, it could easily be adapted to other scientific fields where ultrafast temporal accuracy and signal quality are important.

There are several improvements that can be made to our design. We envision that a future rendition of this device could use fiber optic transmitters and receivers for the output signals to the data recording devices. This would eliminate the need for the opto-isolators, which did introduce some delay into the system. Using fiber optics will also allow further noise reduction, and make cable management easier during experiments as compared to expensive and cumbersome electromagnetically shielded DB25 and DB37 cables. Physical switches for turning off the indicator lights during the experiment could be added to increase the speed and control the room’s lighting during experiments. We considered using the push button function of the rotary encoders to accomplish this, and the code supports this function. However, it is also necessary to consider that adding physical switches in the code can take up processing time.

After our first prototype was put into steady use, we learned that the switch on the battery pack that we chose wore out really quickly. Perhaps a more robust switch for the battery pack, accessible from the outside of the enclosure, would be more suitable. A future version of this project could eliminate the Arduino and have a custom printed circuit board. In addition, a better enclosure material could be chosen. We used Medium Density Fiber board to make our enclosure for ease of prototyping (i.e. Figures 6, 7, and 8). One could use a 3D printer to print the enclosure in plastic. Perhaps the best choice would be to use an electrically isolating material like hard rubber (34).

**Fig. 5.**
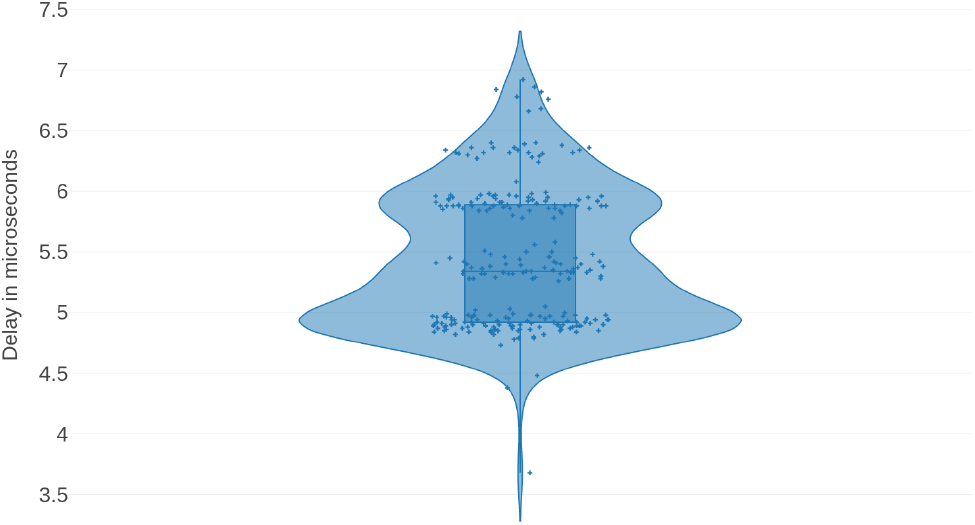
This figure shows the distribution of the delay in microseconds between the input and output of the box for one channel as calculated by the oscilloscope.

**Fig. 6.**
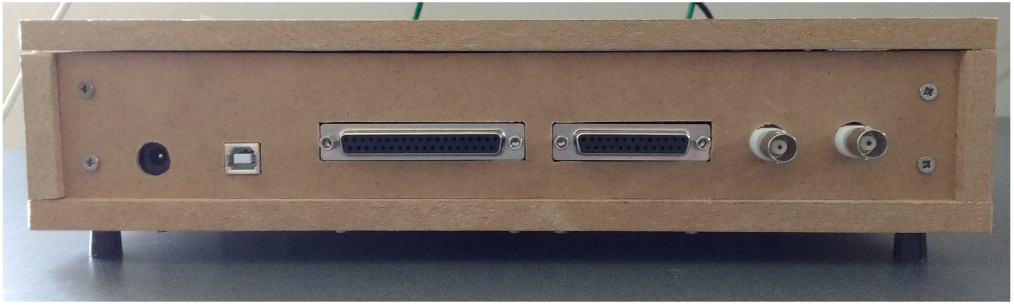
The back panel of the device is shown here. From left to right, the connections are: 2.1mm barrel connector for the power supply, USB port for updating the software, DB37, DB25, photodiode output BNC connector, and microphone output BNC connector.

**Fig. 7.**
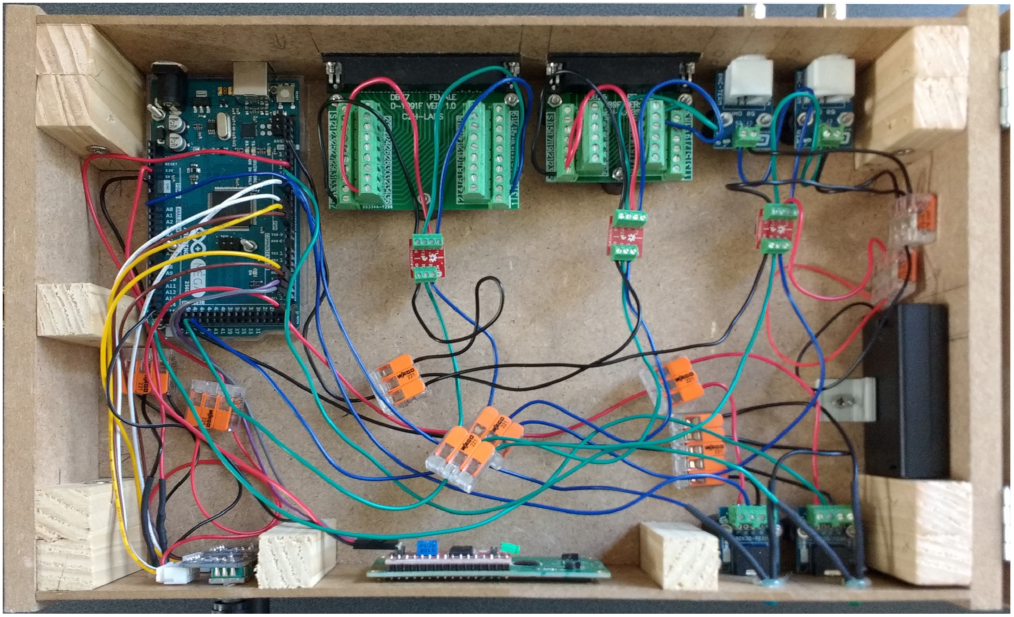
This is an overhead view of the device showcasing the internal wiring. This is a different configuration than that of Figure 3, and is for presentation purposes only. The connections for the rotary encoders and the LCD are different here. To build the device, follow the circuit diagram, Arduino pin map, and code.

**Fig. 8.**
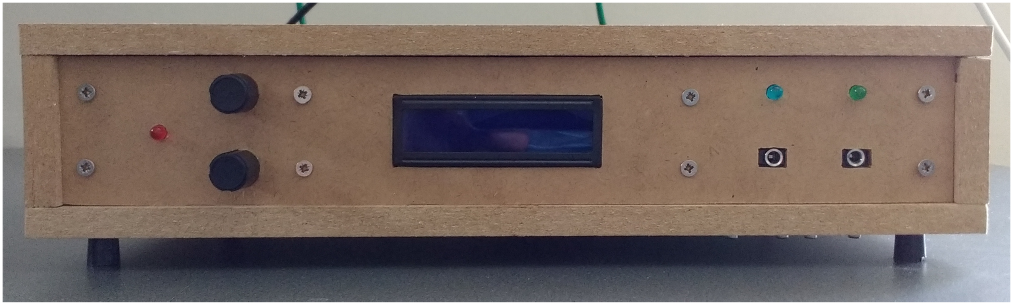
This is the front panel and user interface for the device. From left to right, the elements are: red power indicator light, rotary encoder adjustment knobs, LCD, photodiode and microphone input jacks with corresponding blue and green indicator lights.

Based on this paper, our project is easily reproducible. As we have open-sourced the design and software (29), anyone will be able to improve or use this design however they choose. One could simply use a wiring scheme as described in this paper, or they could have a printed circuit board made once the design is finished. Anyone can make their own or have a professional company make one for them. Maybe, owing to its low cost, open codebase, and opto-isolated and ultra-low latency processing times, the device could provide a verifiable standard for any experimental research project that will be published and requires verification of ultrafast delivery of stimulus onsets to multiple recording systems.

## Acknowledgments

CED and JGM wrote the paper, designed and built the devices, ran tests and experiments, and wrote the software. CED and JGM designed the figures. CED drew the circuit diagrams in Figures 3 and 4, and CED took photographs of the device for Figures 6, 7, and 8. SJT supported the project and helped proofread the paper. The authors would like to thank Dr. Karine Bouyer for her tireless efforts in helping us source all the materials that we needed.

This work received funding from the European Research Council under the European Union’s Seventh Framework Programme (FP/2007-2013) / ERC Grant Agreement n.323711 (M4 project).

